# Concentration-dependent duality of bFGF in regulation of barrier properties of human brain endothelial cells

**DOI:** 10.1101/2021.01.07.425677

**Authors:** Karolina Kriaučiūnaitė, Agnė Pociūtė, Aida Kaušylė, Justina Pajarskienė, Alexei Verkhratsky, Augustas Pivoriūnas

## Abstract

Multiple paracrine factors regulate barrier properties of human brain capillary endothelial cells (BCECs). Understanding precise mode of action of these factors remains a challenging task because of the limited availability of functionally competent BCECs and use of serum-containing medium. In the present study we employed defined protocol for producing BCECs from human inducible pluripotent stem cells. We found that autocrine secretion of basic fibroblast growth factor (bFGF) is necessary for the establishment a tight BCECs barrier, as revealed by measurements of trans-endothelial electric resistance (TEER). In contrast, exogenous bFGF in concentrations exceeding 4 ng/ml inhibited TEER and proliferation of BCECs in a concentration-dependent manner. Exogenous bFGF did not significantly affect expression and distribution of tight junction proteins claudin-5, occludin and ZO-1. Treatment with FGF receptor blocker PD173074 (15 μM) suppressed inhibitory effects of bFGF and induced nuclear translocation of protein ZO-1. Inhibition of phosphoinositide 3-Kinase (PI-3K) with LY294002 (25 μM) significantly potentiated inhibitory effect of bFGF on TEER indicating that PI-3K signalling pathway partially suppress inhibitory effects of bFGF on TEER. In conclusion we show that autocrine bFGF secretion is necessary for the proper barrier function of BCECs, whereas exogenous bFGF suppresses barrier resistance in a concentration-dependent manner. Our findings demonstrate a dual role for bFGF in the regulation of BCEC barrier function.

## Introduction

The blood brain barrier (BBB), formed by the continuous layer of tightly sealed specialised brain capillary endothelial cells (BCECs), separates brain parenchyma from the circulating blood [1]. Endothelial cells together with pericytes, astroglial cells and neurones form the neurogliovascular unit (NGVU) that regulates BBB and local cerebral blood flow [2,3]. At the cellular level, close contacts between BCECs are reinforced by tight and adherens junctions (TJ and AJ respectively) that seal the barrier. The BCECs also contact pericytes with which they share common basement membrane, while at the parenchymal side of the BBB astroglial endfeet plaster the basement membrane of capillaries or parenchymal basement membrane of larger calibre vessels [3]. The TJs and AJs by cementing intercellular contacts between adjacent BCECs prevent paracellular flux of hydrophilic molecules between the blood and the brain. Formation of these intercellular junctions is controlled by multiple paracrine signals deriving mostly from pericytes and astrocytes [4]. The AJs are formed by vascular endothelial (VE) cadherins connecting cytoskeletons of adjacent cells. Under normal conditions cortical actin bundles ascertain equal distribution of AJs between neighbouring cells, whereas permeability increasing factors, such as for example, thrombin, may induce actin remodelling leading to relocation of VE-cadherin complexes and AJs thus disrupting continuity and integrity of BBB [5,6]. The TJ proteins occludin, claudins and zonula occludens (ZO) are crucial regulators of paracellular permeability in the BBB. Cytoplasmic domains of occludin and claudins associate with ZO-1, −2 and −3 proteins, which localise to the cytoplasmic part of the plasma membrane and anchor TJ complexes to the actin filaments [3,5].

Multiple paracrine mechanisms have been implicated in the regulation of expression and recruitment of TJ proteins to the junctional complexes. Astrocytes, for instance, secrete several factors regulating barrier properties of the BBB [7,8]. These astrocytic factors include sonic hedgehog (SHH) [9,10], retinoic acid (RA) [11], ApoE4 and ApoE3 [12], angiopoetins [13], fibroblast growth factor (FGF) [6,14], and glia-derived neurotrophic factor (GDNF) [15] which all contribute to the astrocyte-endothelial, or astrocyte-pericyte-endothelial cells crosstalk. Revealing the mode of action of these paracrine factors is a task far from trivial. The *in vivo* models are not particularly suitable for the precise control and dynamic monitoring of paracrine factors at the BBB. On the other hand, until recently BBB *in vitro* models were limited to either primary, or immortalised BCECs, characterised by low transendothelial electrical resistance (TEER) [16,17]. In addition, most protocols used serum-containing medium, with its undefined components limiting consistency and reproducibility. Both human plasma derived serum and foetal bovine serum (FBS) represent rich and varying source of different growth factors and cytokines that may critically affect BCECs differentiation which further complicate *in vitro* studies of paracrine signalling.

Recent development of fully defined differentiation protocol [18] for producing BCECs from human inducible pluripotent stem cells (iPSCs) allows generation of monocultures with high TEER in the range of ~ 4000 Ω∙cm^2^ that is close to the readings obtained *in vivo* [19]. In the present study we used this protocol to systematically test effects of basic fibroblast growth factor (bFGF) on the barrier properties of human BCECs monolayers. This reductionist approach has enabled identification of a novel role for bFGF in the regulation of barrier properties of BCECs. Autocrine bFGF secretion was necessary for the proper barrier function of BCECs whereas exogenous bFGF in concentrations exceeding 4 ng/ml inhibited TEER in a concentration-dependent manner. Our findings suggest the importance of bFGF signalling in regulation of BBB integrity.

## Materials and methods

### Human exfoliated deciduous teeth stem cells reprogramming into iPSCs

Human exfoliated deciduous teeth stem cells (SHEDs) were derived from milk tooth according to the previously described protocol [20]. Material was collected under the approval of the Lithuanian Bioethics committee. SHEDs were maintained in DMEM-GlutaMAX medium (1 g/L glucose; Gibco), supplemented with 10 % FBS, 100 U/ml penicillin and 100 μg/ml streptomycin (all from Biochrom, Berlin, Germany) at 37 °C in a humidified 5% CO^2^ atmosphere, medium was changed every 2-3 days. iPSCs from SHEDs were generated with Epi5 Episomal iPSC Reprogramming Kit and Neon Transfection System (both from Thermo Fisher Scientific, CA, USA) according to the manufacturer’s protocol. On the 15th day of reprogramming N2B27 medium was switched to complete Essential 8 medium (E8; Thermo Fisher Scientific), which was then changed every two days. By 21st day, when iPSC colonies emerged, they were picked and expanded on plates coated with Matrigel (Corning, Cambridge, United Kingdom). Picked colonies were maintained in complete E8, medium was changed daily.

### iPSC maintenance

In this study we used two iPSC lines. The MBE 2960 line [21] derived from healthy (age at biopsy 78, male) donor was provided by Prof. Alice Pébay, the University of Melbourne and Prof Alex Hewitt, the University of Tasmania. Second iPSC line was derived from SHEDs obtained from healthy human exfoliated deciduous teeth of children (female, 7 years old) at the State Research Institute Centre for Innovative Medicine. iPSCs were maintained in E8 medium supplemented with 50 U/ml penicillin and 50 μg/ml streptomycin, medium was changed every 24 hours. Cells were passaged upon reaching 80% confluence by washing twice with 0.5 mM EDTA solution (Thermo Fisher Scientific), incubating in 0.5 mM EDTA for 3-4 minutes at 37 °C, then detaching by stream of culture medium and subsequent plating onto Matrigel-coated plates in the fresh culture medium, supplemented with 5 μM Y-27632 (Merck). iPSCs used in the following experiments were between passages 15 and 21.

### iPSC differentiation to the brain endothelial cells

Differentiation of iPSCs was based on the recently published protocol [18] with slight modifications. Briefly, iPSCs were seeded on Matrigel-coated plates at a density of 15 800 cells/cm^2^ in E8 medium supplemented with ROCK inhibitor 10 μM Y-27632. Differentiation was initiated after 24 hours by changing medium to Essential 6 (E6; Thermo Fisher Scientific). E6 medium was changed every 24 hours for 4 days. On the 5th day after seeding, medium was changed to human endothelial serum free medium (hESFM); supplemented with 20 ng/ml bFGF (both from Thermo Fisher Scientific), 10 μM retinoic acid (Merck Darmstadt, Germany) and 0.25x B-27 (Thermo Fisher Scientific). After 48 hours the same medium was changed again. The next day cells were subcultured: washed with PBS (Biochrom) once and subsequently incubated with Accutase (BD Biosciences) for 10-20 min. The obtained cell suspension was centrifuged at 300 g for 3 min. Cells were resuspended in hESFM medium supplemented with 0.25x B-27 (Thermo Fisher Scientific), 50 U/ml penicillin and 50 μg/ml streptomycin (further – BCECs culture medium), then seeded onto polyester membrane Transwell inserts (0.33 cm^2^, 0.4 μm pore size; Corning) coated with a mixture of 400 μg/ml collagen IV and 100 μg/ml fibronectin (both from Merck).

### BCEC treatments

1 day after subculturing on Transwell filters medium was changed and various growth factors and pharmacological agents were added (Fig. 1a); after 3 days of further cultivation with these agents the TEER was measured (see Fig. 1a for experimental design). Immediately after these measurements cells were fixed for immunocytochemistry.

**Fig. 1.**
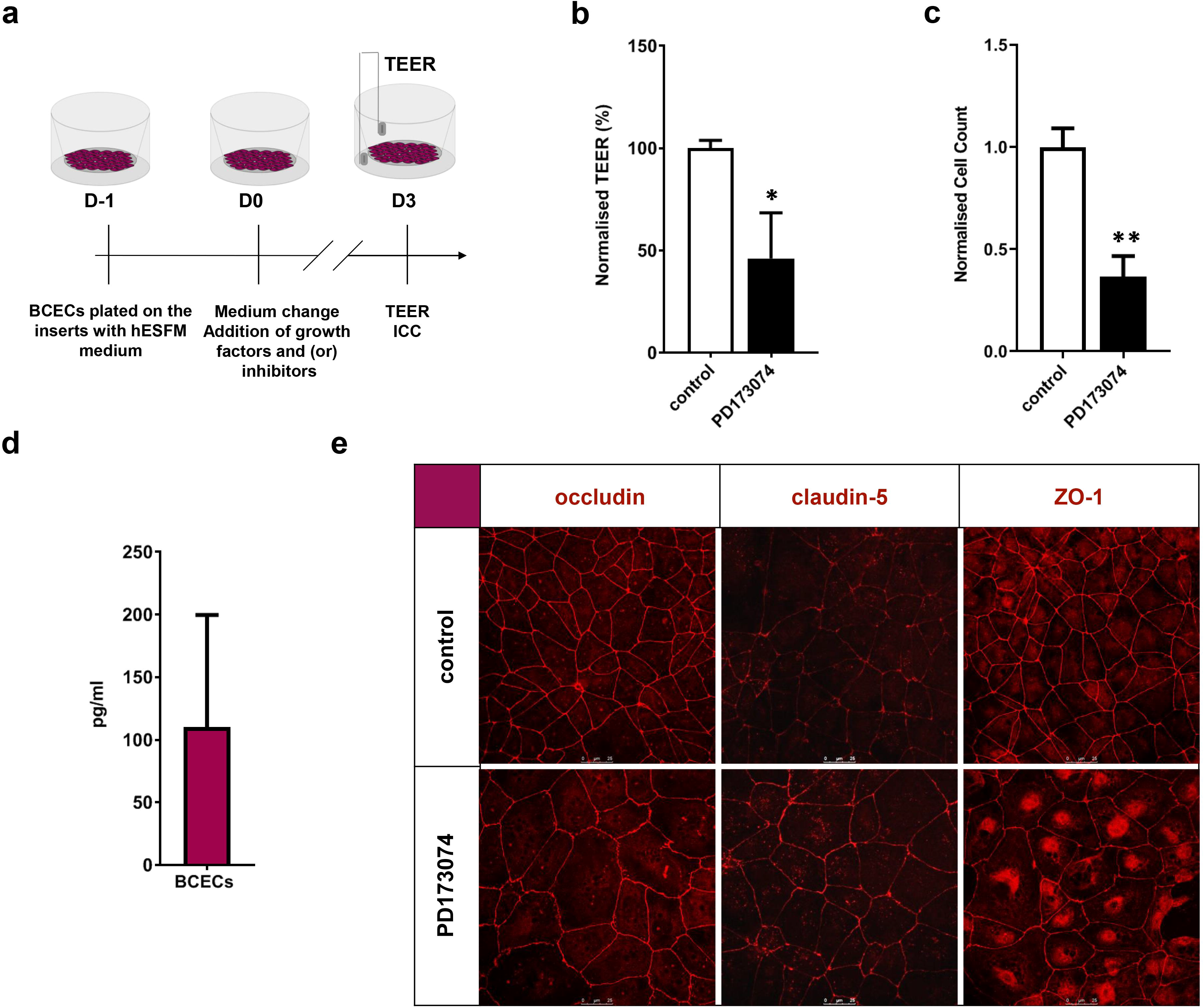
bFGF constitutively secreted by BCECs supports the functional barrier through interacting with TJs and cell proliferation **a** TEER in control conditions and after treatment with selective FGF receptor blocker PD173074 (15 μM) **b** Number of BCECs in control conditions and after treatment with PD173074 (15 μM) **c** bFGF levels measured with ELISA in supernatants from BCECs cultures **d** Representative confocal images of BCEC cultures stained with antibodies against tight junction proteins claudin-5, occludin and ZO-1 Data are normalised to control values and expressed as % ± S.D., * *p* <0.001, ** *p* < 0.0001, *n* = 8

### Transendothelial electrical resistance (TEER) measurements

TEER measurements [22] were performed on 3rd day of incubation with growth factors and (or) different inhibitors by using Millicell ERS-2 Electrical Resistance System (Merck-Millipore). Each insert was measured in three different locations. In order to calculate TEER (Ω·cm^2^), the mean electrical resistance of cell-free insert was subtracted from the mean readings of insert with BCECs and then multiplied by the surface area of the insert.

### Immunocytochemistry and confocal microscopy

After TEER measurements, culture medium was aspirated and cells were fixed with ice cold methanol-acetone solution (1:1) for 10 min at −20 °C, washed three times with PBS and blocked using 1 % BSA-PBS solution for 30 min at room temperature (RT). Then incubated with primary antibodies (diluted in 1 % BSA-PBS) against ZO-1 (1:33), claudin-5 (1:100) and occludin (1:50) (all from Thermo Fisher Scientific) at 4 °C overnight. Next, cells were washed three times with PBS and incubated with secondary antibodies diluted in PBS (1:1000 Alexa Fluor 594, Thermo Fisher Scientific) for 1 hour at RT, in the dark. Alternatively, cells were fixed with 4% PFA for 20 min at RT, washed three times with PBS, permeabilised with 0.1% Triton X-100 for 15 min, washed three times with PBS again, blocked and incubated with Alexa Fluor 647-phalloidin conjugate (1:40; Life Technologies) for 1 h at RT in the dark. Then all membranes were washed 3 times and mounted on coverslips using an aqueous fluorescent mounting medium (DakoCytomation, Huddinge, Sweden). Samples were analysed with Leica TCS SP8 confocal microscope (Leica Microsystems, Mannheim, Germany), using Diode 405 nm, DPSS 561 nm and a HeNe 633 nm lasers. Images were taken using 63x oil immersion lens.

### ELISA

The levels of bFGF autocrine secretion in BCECs monolayers were evaluated after 3 day incubation in hESFM medium. Cell culture supernatants were centrifuged at 10 000 g for 10 min and stored −20 °C until further analysis. The levels of bFGF were quantified with human bFGF ELISA kit (BioLegend, Cat. Nr. 434309) according to the manufacturer’s instructions. The colorimetric measurements (λ=450 nm, reference λ=570 nm) were performed using Asys UVM340 plate reader (Biochrom).

### Cell counting

Images taken with confocal microscope were used for cell counting, which was performed manually using ImageJ program, multi-point tool. Only cell nuclei that entirely fit into a field of 185 x 185 μm were counted.

### Statistics

Differences between 2 groups were compared by Student’s t-test. Data from all remaining experiments were compared by one-way ANOVA following Tukey’s or Dunnett’s post-test. All results were considered significant, at *p* < 0.05. Statistical analysis was performed with Graph Pad Prism® software version 8.0.2 (Graph Pad Software, Inc., City, State, USA).

## Results

### Autocrine/paracrine signalling of bFGF is important for the maintenance of proper barrier function in BCECs

After growing on the membraneous inserts for 3 days, the BCECs developed a functional barrier, as confirmed by TEER measurements: the TEER readouts ranged 3582 ± 756 Ω*cm^2^ (n = 19), which compares well with the *in vivo* readouts of ~ 5000 – 6000 Ω*cm^2^ [19]. The BCECs monolayers were also characterised by membrane-associated expression of major TJ proteins as revealed by immunochemistry (Fig. 1). Treatment of BCECs monolayers with selective inhibitor of bFGF receptor PD173074 (15 μM) resulted in a significant decrease (by 53.83 ± 22.28 %; n = 8) of TEER values (Fig. 1b). This decrease coincided with substantial reduction in the number of BCECs in monolayers (by 63.37 ± 10.07 %, n = 8, Fig. 1c). These effects of PD173074 suggest constitutive secretion of bFGF which regulates barrier function through autocrine/paracrine stimulation of bFGF receptors. Indeed, ELISA assays confirmed this assumption by revealing bFGF in the culture media (Fig. 1d). At a sub-cellular level, inhibition of bFGF receptors with PD173074 induced redistribution of ZO-1 towards the cell nuclei; the immunolocalisation of claudin-5 and occludin, however, was not affected (Fig. 1e).

### Exogenous bFGF inhibits TEER and increases BCEC proliferation in a concentration-dependent manner

Treatment of BCECs cultures with exogenous bFGF affected TEER in the concentration-dependent manner. At the lowest concentration of 0.8 ng/ml no changes in TEER were detected, however at higher concentrations TEER was progressively decreased (Fig. 2a). At 4 ng/ml bFGF the TEER was decreased by 65.29 ± 6.75 %, at 8 ng/ml by 84.96 ± 5.13 and at 16 ng/ml by 97.06 ± 1.55% (n = 6); the apparent EC_50_ being 3.2 ng/ml. At the same time treatment of BCECs cultures with increasing doses of bFGF elevated number of cells in a similar concentration-dependent fashion (Fig. 2b). At 0.8 ng/ml bFGF increased cell counts by 32.70 ± 12.29 % (n = 6), at 4 ng/ml by 73.30 ± 22.56 % (n = 6), at 8 ng/ml by 95.00 ± 14.64 % (n = 6), and at bFGF 16 ng/ml by 99.00 ± 16.06 %, (n = 4); with the apparent EC_50_ of 2.2 ng/ml.

**Fig. 2.**
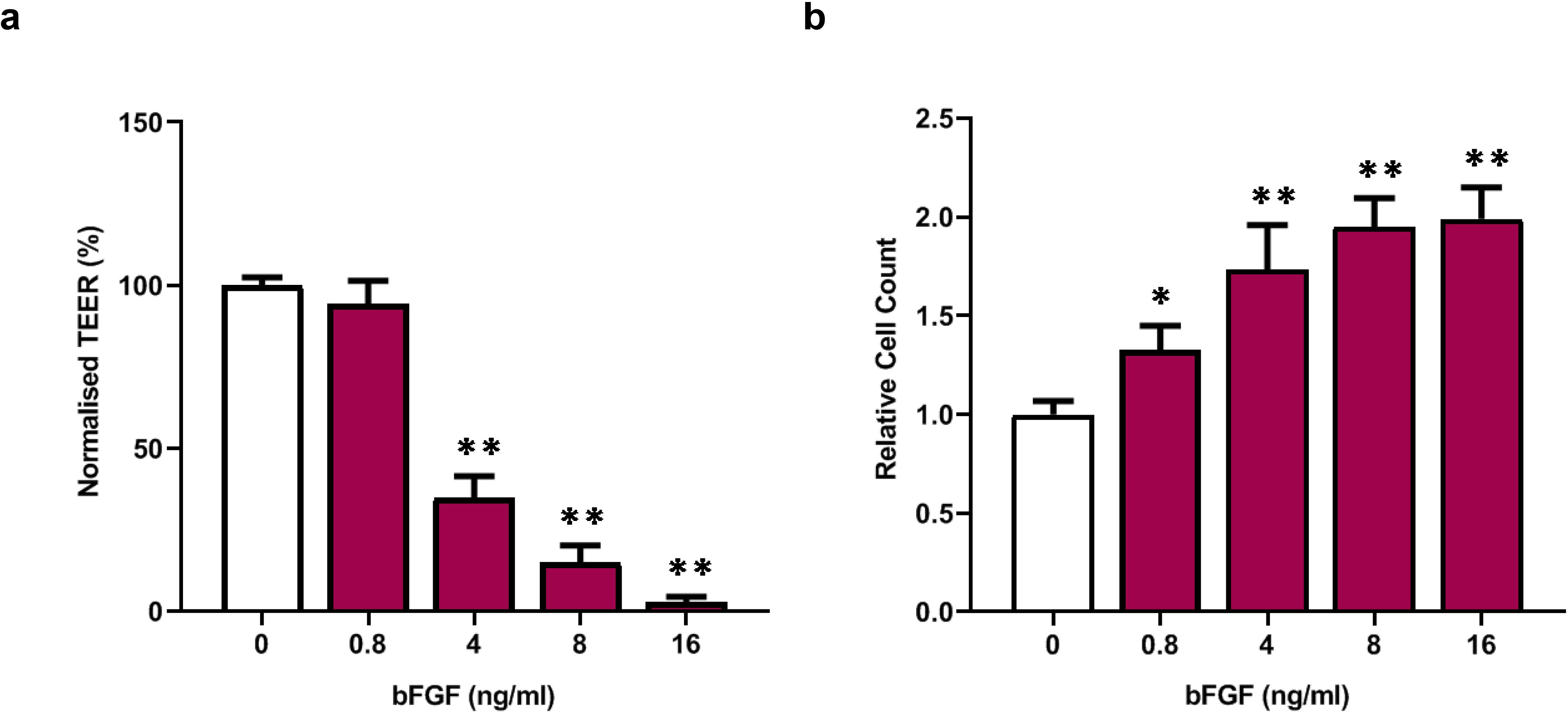
bFGF decreases TEER and increases number of BCECs in a concentration-dependent manner **a** TEER in control cultures and after treatment with different concentrations of bFGF (0.8, 4, 8 or 16 ng/ml) **b** Number of BCECs in control conditions and after treatment with different concentrations of bFGF Data are normalised to a control values and expressed as % ± S.D., * *p* < 0.001, ** *p* < 0.0001, *n* = 4-6

### Specific inhibitor of bFGF receptor PD173074 antagonises effects of exogenous bFGF

Treatment of BCECs with bFGF (8 ng/ml) suppressed TEER by 74.12 ± 26.95 % (n = 9), whereas combined treatment with bFGF and inhibitor of bFGF receptor PD173074 (15 μM) suppressed TEER only by 39.53 ± 21.55 % n = 9, indicating that PD173074 partially antagonised inhibitory effect of bFGF on the TEER readouts (Fig. 3a). Treatment of BCECs with bFGF (8 ng/ml) increased cell number by 95.00 ± 15.86 % (n = 9), whereas combined treatment with PD173074 decreased cell counts by 64.71 ± 8.06 % (n = 7) (Fig. 3b). Treatment of BCECs cultures with combination of bFGF and PD173074 affected neither expression nor distribution of claudin-5 and occludin (Fig. 3c). Exogenous bFGF also did not affect PD173074-induced nuclear accumulation of ZO-1 (Fig. 3c).

**Fig. 3.**
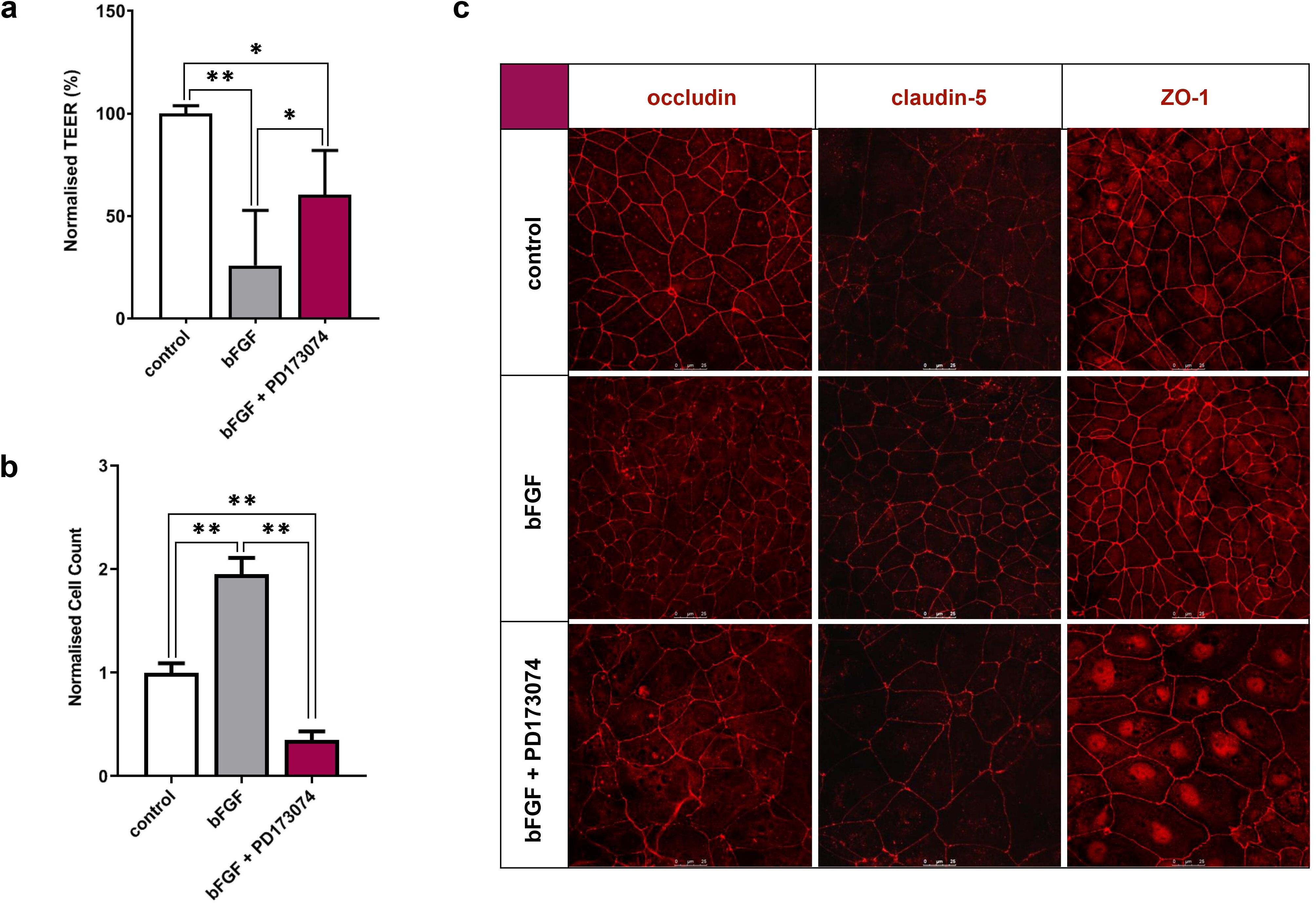
Selective FGF receptor inhibitor PD173074 antagonises effects of exogenous bFGF **a** TEER in control cultures and after treatment with bFGF (8 ng/ml) alone or in combination with PD173074 (15 μM) **b** Number of BCECs in control cultures and after treatment with bFGF (8 ng/ml) alone or in combination with PD173074 (15 μM) **d** Representative confocal images of BCECs cultures stained with antibodies against tight junction proteins claudin-5, occludin and ZO-1 Data are normalised to control values and expressed as % ± S.D., * *p* < 0.001, ** *p* < 0.0001, *n* = 7-9

### Effects of different combinations of bFGF, EGF, BMP-2 and CNTF on the barrier properties of BCECs

We next investigated effects of bFGF in combination with different growth factors (GFs) on TEER in BCECs monolayers (Fig. 4). Treatment with combination of GFs consisting of bFGF (8 ng/ml), EGF (10 ng/ml), BMP-2 (10 ng/ml) and CNTF (5 ng/ml) significantly decreased TEER values in BCECs monolayers (Fig.4). After 3 days of incubation with GFs, TEER decreased up to 13.9 times when compared to untreated BCECs. All GF combinations containing bFGF strongly inhibited TEER (Fig. 4). The EGF potentiated inhibitory effect of bFGF, while EGF alone, or in combinations with other GFs decreased TEER only moderately. We also found that treatment with BMP-2, or CNTF alone as well as in combination, significantly increased TEER values in BCECs monolayers. Thus, bFGF strongly inhibits TEER also in the presence of other GFs.

**Fig. 4.**
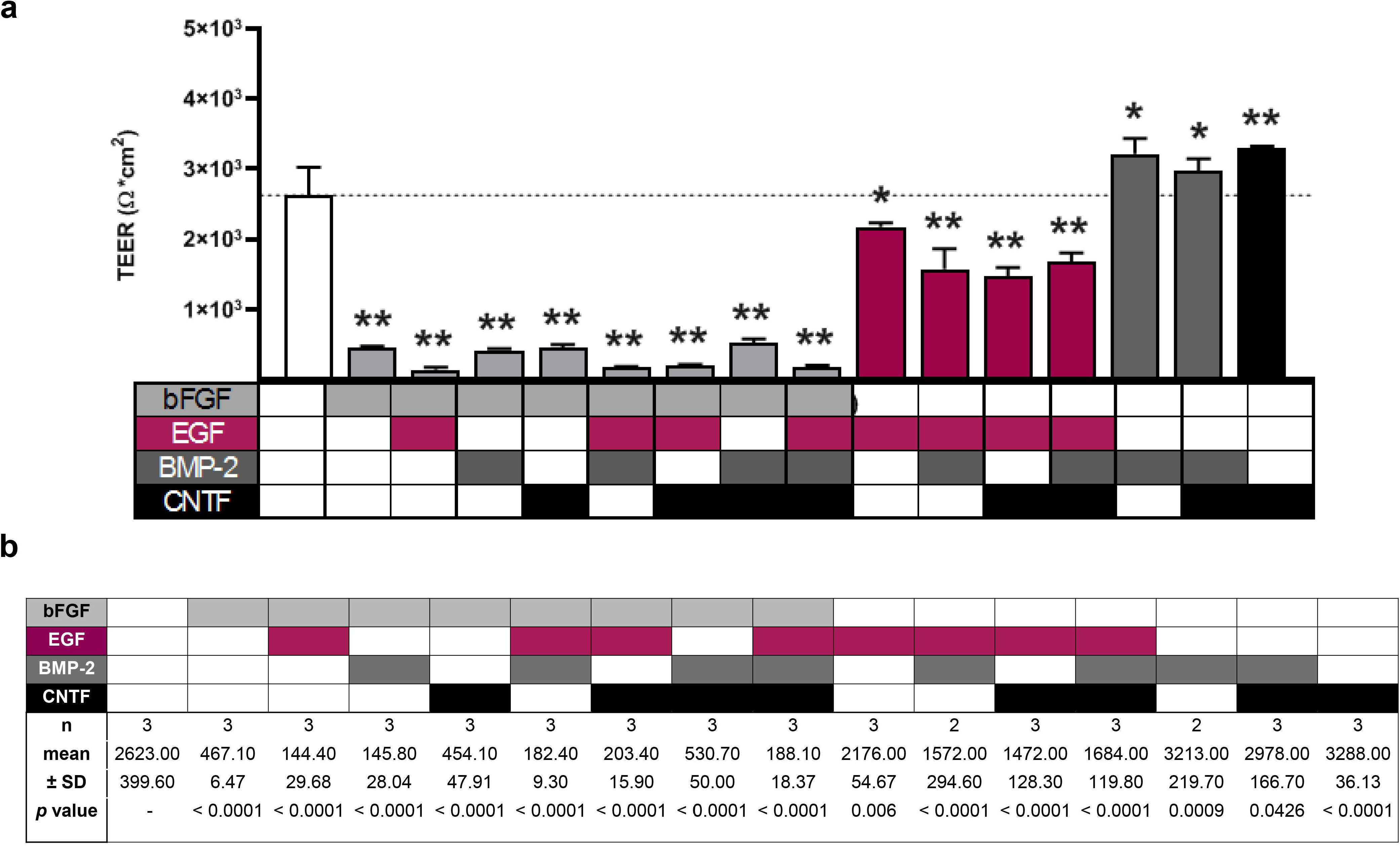
TEER measurements in the presence of several GFs **a** TEER readouts after treatment with bFGF (8 ng/ml), EGF (10 ng/ml), BMP-2 (10 ng/ml) and CNTF (5 ng/ml) alone or in combinations **b** Table of descriptive statistics TEER is expressed as Ω*cm^2^ ± S.D., * *p* < 0.05, ** *p* < 0.0001, *n* = 2-3 inserts

### Inhibition of phosphoinositide 3-Kinase (PI-3K) potentiate inhibitory effect of bFGF on TEER

Treatment of BCEC monolayers with PI-3K inhibitor LY294002 (25 μM) alone down-regulated TEER by 47.63 ± 23.53 % (n =8) (Fig. 5a), while bFGF alone down-regulated TEER by 74.43 ± 5.84 % (n = 8) (Fig. 5a). Combination of bFGF (8 ng/ml) with LY294002 potentiated action of bFGF on TEER, decreasing it to 99.22 ± 0.33 % when compared to control (n = 8). Exposure to LY294002 inhibited proliferation of bFGF-treated and untreated BCECs by 32.50 ± 19.69 % (n = 8) and by 47.44 ± 10.26 %, (n = 9), respectively (Fig. 5b). Immunocytochemistry revealed fragmented staining patterns of occludin and ZO-1 of BCECs treated with bFGF and LY294002 (Fig. 5c).

**Fig. 5.**
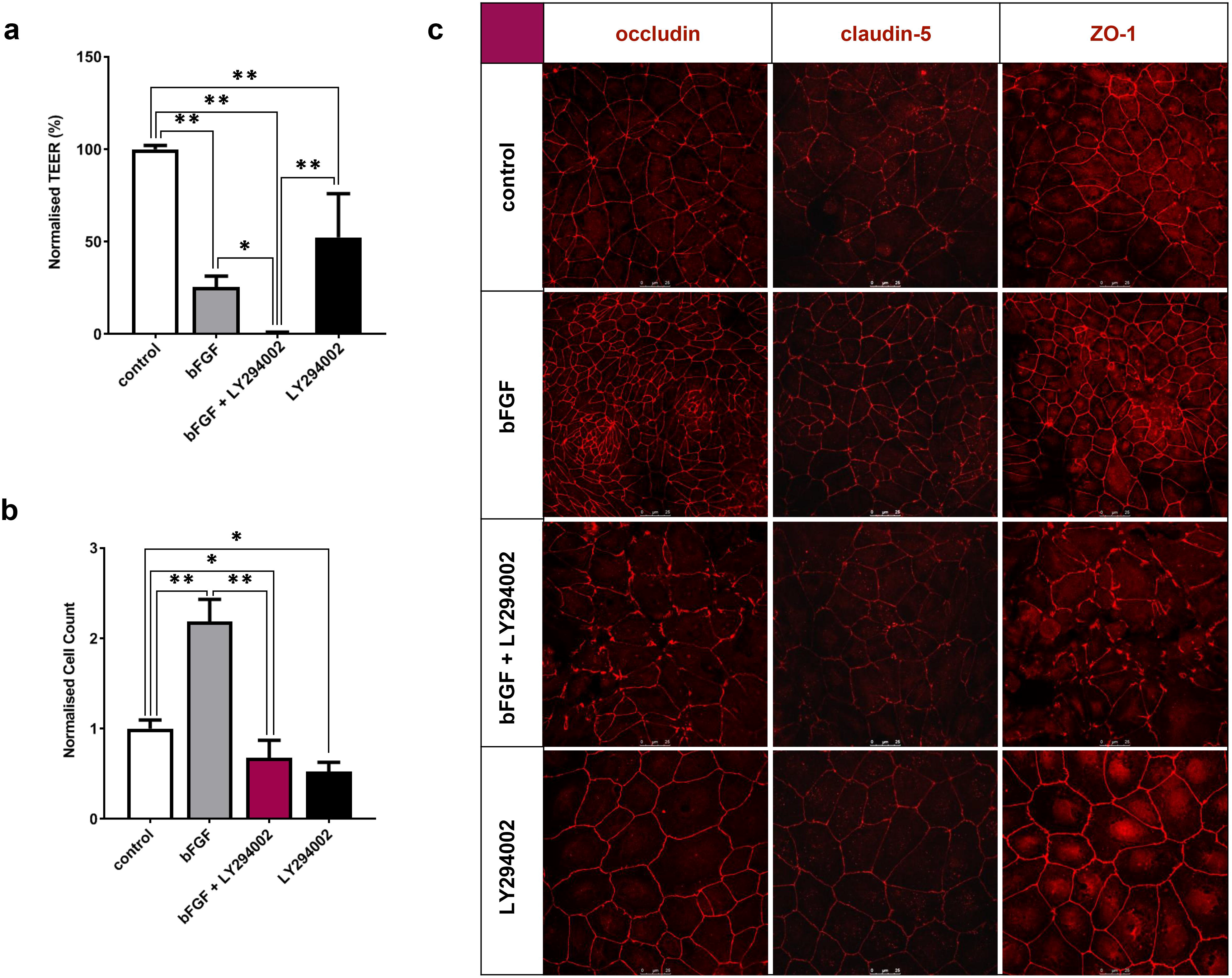
PI 3-kinase inhibitor LY294002 decreases TEER andpotentiates bFGF effects **a** TEER readouts after treatment with bFGF (8ng/ml) alone or in combination with LY294002 **b** Number of BCECs in control conditions and after treatment with bFGF (8ng/ml) alone or in combination with LY294002 **c** Representative confocal microscopy images of BCECs cultures stained with antibodies against tight junction proteins claudin-5, occludin and ZO-1 Data are normalised to control values and expressed as % ± S.D., * *p* < 0.001, ** *p* < 0.0001, *n* = 6-9

Treatment with ROCK inhibitor Y-27632 (10 μM) alone or in combination with bFGF did not significantly change TEER in BCEC monolayers. Neither it did affect proliferation (Fig. 6a,b);As expected, staining with phalloidin antibodies showed that blocking of ROCK significantly decreased formation of stress fibres in control and bFGF – treated cells (Fig. 7).

**Fig. 6.**
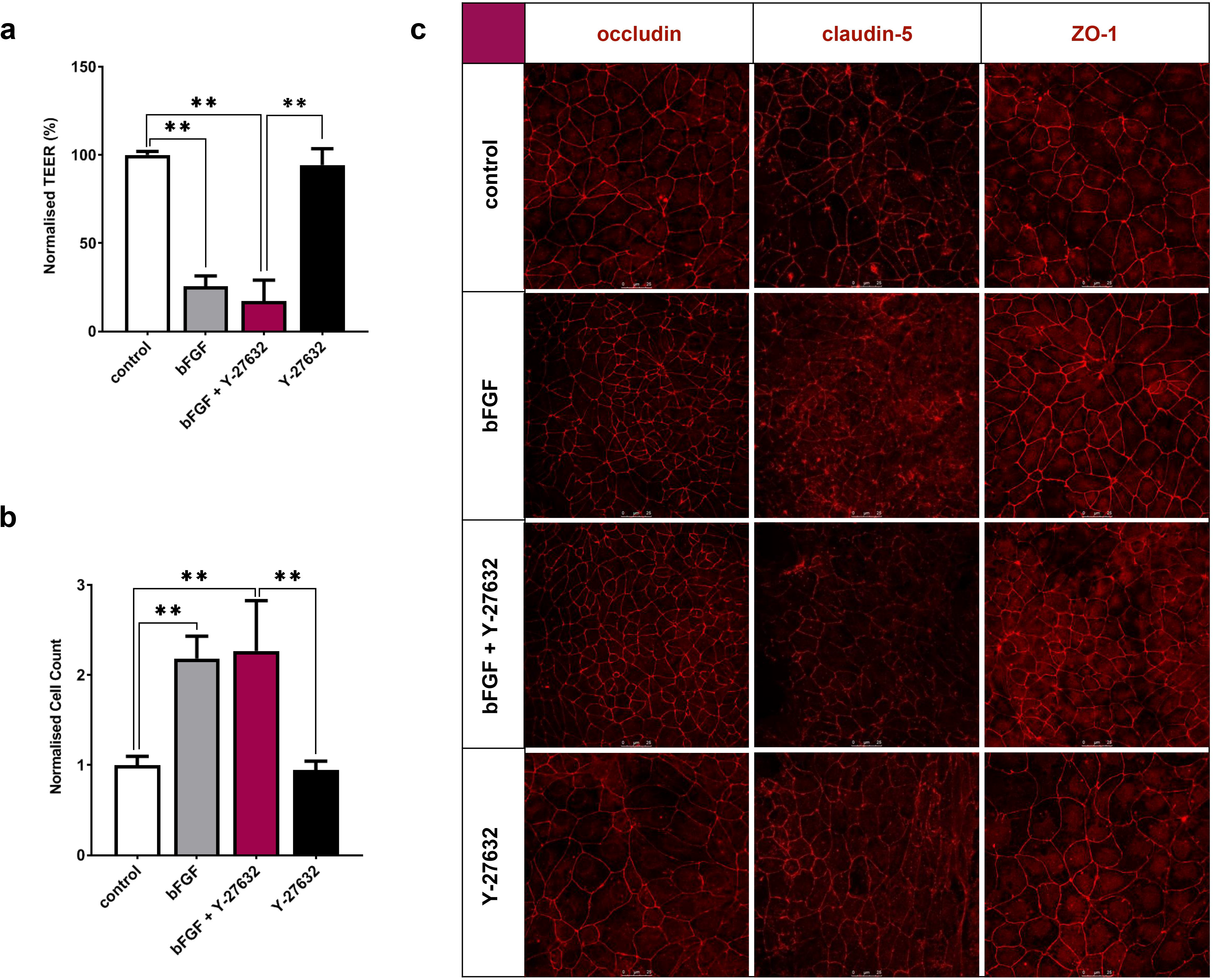
ROCK inhibitor Y-27632 strengthened bFGF effect on TEER in BCECs, but had no effect by itself **a** TEER readouts in control conditions and after treatment with bFGF (8ng/ml) alone or in combination with Y-27632 **b** Number of BCECs in control conditions and after treatment with bFGF (8ng/ml) alone or in combination with Y-27632 **c** Representative confocal images of BCEC cultures stained with antibodies against tight junction proteins claudin-5, occludin and ZO-1 Data are normalised to control values and expressed as % ± S.D., * *p* < 0.001, ** *p* < 0.0001, *n* = 6-8

**Fig. 7.**
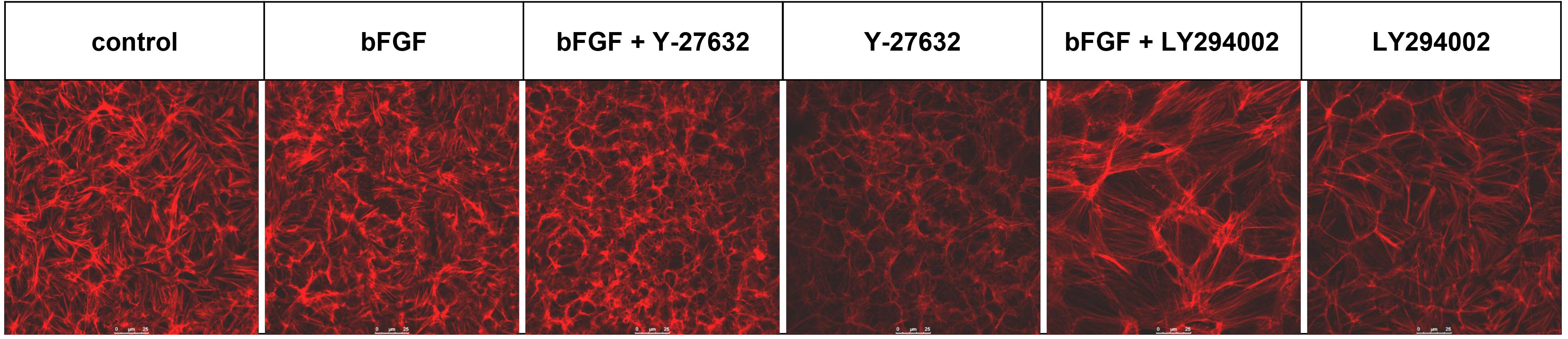
ROCK inhibitor Y-27632 reduced stress fibre formation in untreated and bFGF-treated BCECs Representative confocal microscopy images of BCEC cultures stained with antibodies against F-actin.

## Discussion

In the present study we, for the first time, demonstrate a dose and source dependent duality of bFGF in the regulation of BCECs barrier integrity. First, we show that autocrine secretion of bFGF by BCECs is necessary for the proper barrier function; this action is achieved at low (< 4 ng/ml) concentration of bFGF. Second, we demonstrate that exogenous bFGF in concentrations above 4 ng/ml effectively suppresses TEER in BCEC monolayers (Fig. 8). Our findings suggest a novel mechanism for the effects of bFGF on the BCEC barrier function with potential implications in both physiological and pathological conditions.

**Fig. 8.**
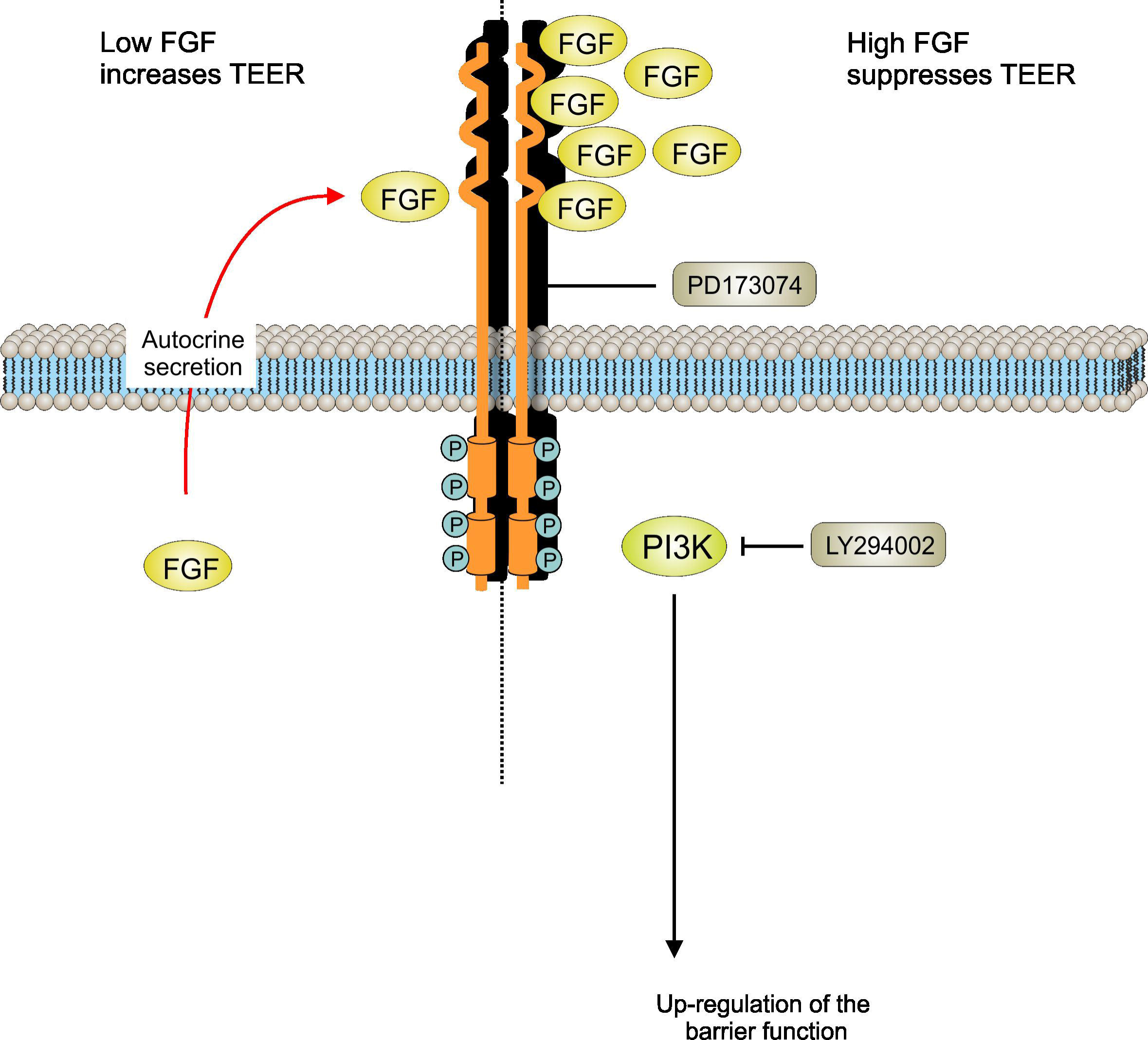
Dual action of bFGF on the brain endothelial cells barrier. See text for explanation

The bFGF exerts pleiotropic effects in various tissues and acts as an important modulator of angiogenesis [23], CNS development [24], adult neurogenesis and neuroinflammation [25]. Several studies investigated effects of exogenous bFGF administration on the BBB integrity. All reports published hitherto demonstrated protective role of exogenous bFGF against BBB disruption occurring after intracerebral haemorrhage [6], traumatic brain injury [26], oxygen glucose deprivation [27,28] and mannitol-induced hyperosmotic shock [29]. In consequence, it was proposed that bFGF preserves AJs by suppressing the RhoA and ROCK by FGFR-induced activation of the PI3K-Akt-Rac1 signalling pathway [6]. Several *in vitro* studies [27,29] confirmed protective role of bFGF on the TEER of BCECs and are therefore in disagreement with our data. These discrepancies likely reflect differences in cell identities and in experimental design. More specifically, high concentration of bFGF (2.5 μM, or ~ 42.5 μg/ml) has been used prior to TEER recordings in primary human brain microvascular endothelial cells (hBMEC) [27]. The study in question is also vague about how bFGF or PD173074 affected TEER values in hBMEC monolayers. Another investigation [30] used BCECs differentiated from human iPSCs according to the similar protocol as used by us. However, this protocol included platelet-poor plasma-derived serum which introduces an unknown and varying mixture of different growth factors; in contrast we applied a serum-free protocol [18].

We demonstrate that differentiated BCECs constitutively secrete bFGF and that autocrine/paracrine bFGF signalling is important for the maintenance of proper barrier resistance: inhibition of bFGF receptors with specific blocker leads to a significant disruption of barrier integrity reflected by a substantial (~ 54 %) decrease in TEER values. Besides TEER maintenance, autocrine/paracrine bFGF signalling also positively regulates BCEC proliferation. Treatment with FGF receptor blocker PD173074 induced nuclear translocation of TJ protein ZO-1. In addition to being components of TJs and AJs, ZO proteins can shuttle between the cytoplasm and the nucleus and are involved in signal transduction and regulation of cell growth and proliferation [31]. Several reports demonstrated that nuclear accumulation of ZO-1 is inversely related to the maturity and integrity of cell-cell contacts [32,33]. These observations inspired the assumption that nuclear accumulation of ZO proteins represents a general response of epithelial and endothelial cells to different types of stress [31]. Our findings show similar redistribution of ZO-1 after inhibition of bFGF receptors in BCECs. We therefore suggest that inhibition of bFGF receptor decreases TEER at least partially through relocation of ZO-1 from TJs and AJs to the nuclei of BCECs.

Previous reports demonstrated that bFGF regulates BBB integrity through the PI-3K and ROCK signalling pathways [6,26]. Our data similarly indicate that PI-3K signalling contributes to regulation of BCEC barrier and BCEC proliferation (Fig. 8). Treatment with PI-3K blocker LY29402 significantly potentiated inhibitory effect of bFGF on TEER. It has been shown that PI-3K signalling improves vascular permeability through Akt-FoxO pathway and subsequent upregulation of claudin-5 [34,35]. Akt is also a negative regulator of FoxO1 which can act as a transcriptional repressor of occludin [36]. Immunocytochemical labelling revealed fragmented staining patterns for occludin and ZO-1 in the membranes of BCECs treated with bFGF and LY29402 (Fig. 5c). LY294002 also induced moderate nuclear accumulation of ZO-1. Our results show that PI-3K signalling pathway partially antagonises inhibitory effects of bFGF on TEER.

ROCK promotes breakdown of intercellular junctions by inducing actomyosin contractility and relocation of AJs [5,6]. We demonstrate that inhibiting of ROCK with Y-27632 alone or in combination with bFGF affects neither BCEC proliferation nor TEER. We therefore conclude that in our experimental model ROCK signalling is not involved in the regulation of FGF-dependent effects on BCECs. Besides PI-3K-Akt, activated FGF receptor is coupled to RAS-MAPK, PLCγ and STAT intracellular signalling pathways [37] that can mediate effects of bFGF.

Several studies demonstrated that bFGF can produce opposite effects depending on the concentration. Thus, bFGF can function as either a positive or a negative factor regulating *in vitro* adipogenesis by controlling activation of the ERK signalling pathway [38]. Another report demonstrated that, depending on bFGF concentration, non-canonical TGFβ can either restrict or promote FGFR signalling through ERK-dependent phosphorylation of adaptor protein FRS2 [39]. Further studies are necessary to elucidate detailed mechanisms by which different amounts of bFGF elicit opposite effects on the barrier properties of BCECs.

It is well known that injury and inflammation promote FGF-dependent angiogenesis [23,40]. We suggest that local fluctuations of bFGF may critically affect BBB integrity and permeability *in vivo.* What could be the main cellular source for the bFGF at the brain? Pericytes make close contacts and share common basement membrane with BCECs and therefore could represent potential source for exogenous bFGF. Indeed, pericytes strongly express bFGF and FGFR1 in peri-infarct areas after ischemic insult in mice [41]. Another report demonstrated that peripheral nerve pericytes modify BBB function and TJs partially through the secretion of bFGF [42]. Astrocytes represent another potential source of bFGF [43]. Different CNS pathologies have been associated with increased astroglial expression of bFGF [44]. Thus, acute stress or corticosterone administration induced bFGF secretion in hippocampal astrocytes of rats [45]. Astrocytes associated with white matter expressed high levels of bFGF during initial phase of remyelination after spinal cord lesions [46]. Of course, the results obtained in our *in vitro* study cannot be mechanically extrapolated to the complex *in vivo* contexts. However, they can be used as a starting point for more in-depth studies focusing on simultaneous *in vivo* monitoring of BBB permeability and local expression/secretion of bFGF during various pathologies.

In conclusion we show that autocrine bFGF secretion is necessary for the proper barrier function of BCECs whereas exogenous bFGF suppress it in a dose-dependent manner. Our findings demonstrate a dual role of bFGF in the regulation of BCEC barrier function.

## Acknowledgements

This work was supported by the Global Grant measure (No. 09.3.3-LMT-K-712-01-0082).

## Notes

### Competing Interest Statement

The authors have declared no competing interest.

